# Dysregulation of cell state dynamics during early stages of serous endometrial carcinogenesis

**DOI:** 10.1101/2024.03.15.585274

**Authors:** Andrea Flesken-Nikitin, Matalin G. Pirtz, Christopher S. Ashe, Lora H. Ellenson, Benjamin D. Cosgrove, Alexander Yu. Nikitin

## Abstract

Serous endometrial carcinoma (SEC) constitutes about 10% of endometrial carcinomas and is one of the most aggressive and lethal types of uterine cancer. Due to the rapid progression of SEC, early detection of this disease is of utmost importance. However, molecular and cellular dynamics during the pre-dysplastic stage of this disease remain largely unknown. Here, we provide a comprehensive census of cell types and their states for normal, pre-dysplastic, and dysplastic endometrium in a mouse model of SEC. This model is associated with inactivation of tumor suppressor genes *Trp53* and *Rb1*, whose pathways are altered frequently in SEC. We report that pre-dysplastic changes are characterized by an expanded and increasingly diverse immature luminal epithelial cell populations. Consistent with transcriptome changes, cells expressing the luminal epithelial marker TROP2 begin to substitute FOXA2+ cells in the glandular epithelium. These changes are associated with a reduction in number and strength of predicted interactions between epithelial and stromal endometrial cells. By using a multi-level approach combining single-cell and spatial transcriptomics paired with screening for clinically relevant genes in human endometrial carcinoma, we identified a panel of 44 genes suitable for further testing of their validity as early diagnostic and prognostic markers. Among these genes are known markers of human SEC, such as C*DKN2A,* and novel markers, such as *OAS2 and OASL,* members of 2-5A synthetase family that is essential for the innate immune response. In summary, our results suggest an important role of the luminal epithelium in SEC pathogenesis, highlight aberrant cell-cell interactions in pre-dysplastic stages, and provide a new platform for comparative identification and characterization of novel, clinically relevant prognostic and diagnostic markers and potential therapeutic modalities.

## Introduction

Uterine cancers are projected to be the fourth most common cancer in women in the United States in 2024, and their incidence is increasing about 1-2% per year^1^. While most other types of cancers have a mortality rate trending downwards due to innovations in diagnostic and treatment capabilities, uterine cancer is the only cancer marked by decreased survival over the past 40 years^2^. Endometrioid and serous endometrial carcinomas are the most common subtypes of uterine cancer. Endometrioid endometrial carcinoma constitutes approximately 80% of all endometrial carcinoma cases. With rare exception, these tumors are low grade and have a relatively indolent progression^3,4^. Conversely, serous endometrial carcinoma (SEC) often presents at high stage and is responsible for 40% of uterine cancer related deaths while only constituting about 10% of uterine cancers^4,5^.

If caught early (within stage I or II), SEC often has a favorable five-year survival rate of 74%^6^. However, the survival rate drops rapidly when identified at stage III or IV to about 33%, and most cases aren’t identified until this stage^6^. Another factor affecting the mortality rate of SEC is the high rate of recurrence in patients, which also increases significantly with stage of diagnosis^7^. Thus, there is a significant need to improve the early diagnostic modalities and treatment approaches for SEC patients.

Identification and characterization of early molecular and cellular changes during neoplastic progression are essential for improving cancer detection and treatment. For example, markers of early detection have been explored in breast^8^, ovarian^9^, colorectal^10^, and lung^11^ cancers. Molecular changes in early endometrial carcinogenesis have also been explored^12–15^. However, very few studies focused on SEC, and no new, definitive markers have been identified in the putative SEC precursor lesions, endometrial glandular dysplasia^12–15^. Rapid onset of disease represents a significant challenge for reliable identification of early, pre-dysplastic molecular changes during SEC formation.

Recently, we have established a genetically modified mouse model of SEC^16^. This model is based on an endometrial epithelium-specific conditional inactivation of tumor suppressor genes, *Trp53* and *Rb1*, whose pathways are frequently altered in human SEC^4,17^. In close similarity to their human counterparts, mouse SEC is marked by cytoplasmic expression of p16 (also known as CDKN2A) and reduced expression of progesterone receptor (also known as PGR). The earliest dysplastic lesions are observed 60 days post-induction (DPI) followed by development of overt SEC characterized by invasion into the myometrium and serosa in 81% of mice within the next 340 days^16^.

In the current study, we have investigated single-cell and spatial RNA transcriptome changes in both pre-dysplastic (30-50 DPI) and dysplastic (80-150 DPI) stages of mouse SEC and assessed clinical relevance of our findings to human SEC.

## Results

### Experimental design

We have compiled single-cell and spatial RNA transcriptomes of the mouse endometrium before and after induction of serous endometrial carcinoma (SEC) using our previously described mouse model^16^ (**Figure 1A**). To prepare a census of single-cell RNA expression profiles, we collected uterine horns from 18 mice including 5 normal, 7 pre-dysplastic (30-50 days post-induction, DPI), and 6 dysplastic (80-150 DPI) samples (**Figure 1B**). For spatial transcriptome evaluation we collected uterine horns from 13 mice including 4 normal, 4 pre-dysplastic, and 5 dysplastic samples. All mice were at late diestrus according to vaginal smears followed by morphological verification and estimation of Ki67 index in paraffin sections.

**Figure 1.**
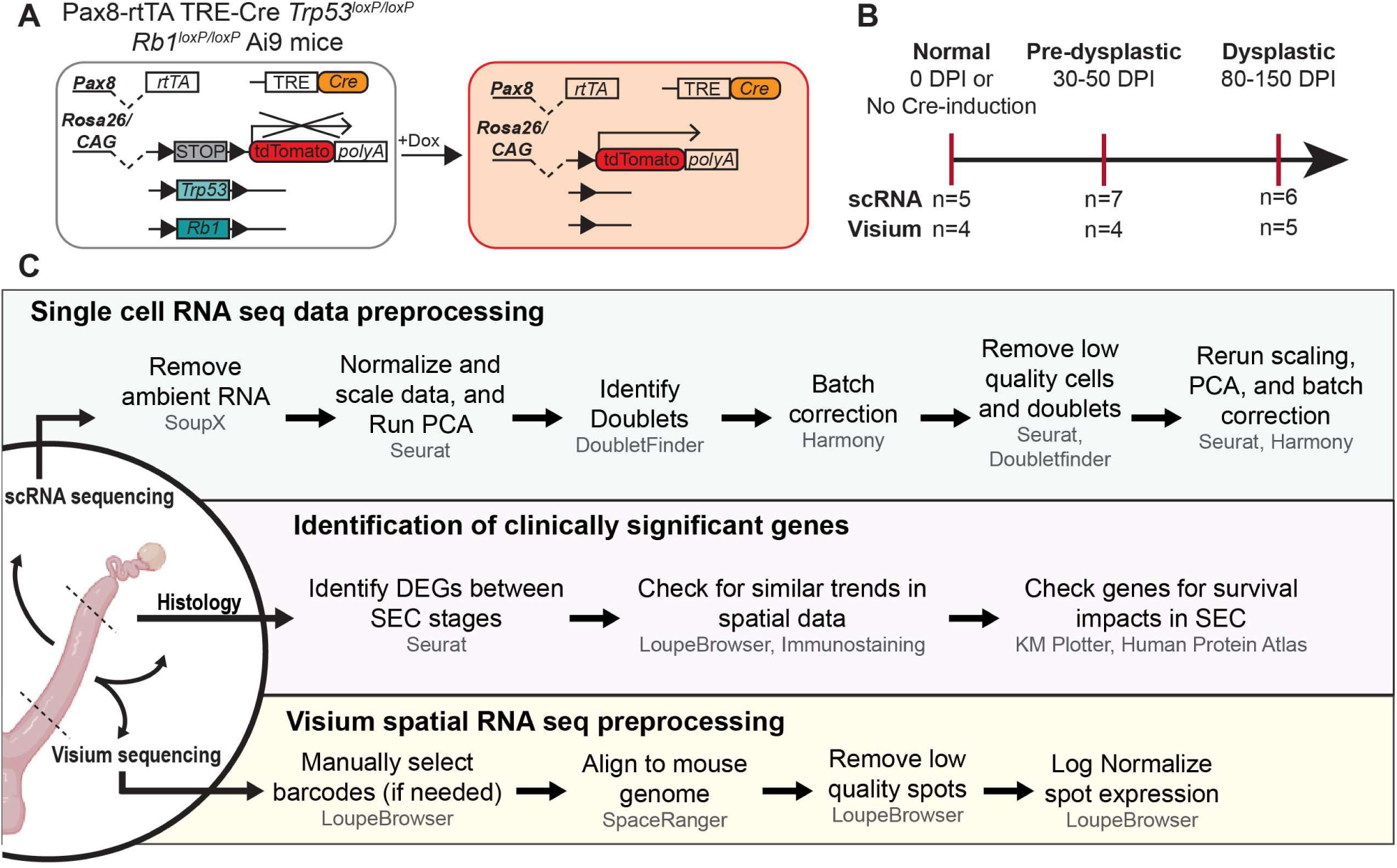
Experimental design. (A) A mouse model of SEC based on conditional inactivation of *Trp53* and *Rb1* in *Pax8*-expressing cells^16^. In this model, the Cre recombinase is activated by doxycycline (Dox). Cre then targets the *loxP* sites flanking exons of the *Trp53* and *Rb1* genes and the STOP codon upstream of the *TdTomato* reporter gene (Ai9), initiating double strand excision of these elements. (B) Data collection timeline. Samples were collected at specific days post induction (DPI) with a single intraperitoneal doxycycline (Dox) injection, for either single cell RNA sequencing or Visium RNA sequencing. n, number of samples per time point. (C) Data processing workflow. Top panel, single cell RNA sequencing preprocessing steps after sample alignment to mouse genome. Middle panel, process by which clinically significant genes were identified. Bottom panel, Visium spatial sequencing preprocessing steps. DEG, differentially expressed genes; PCA, principal component analysis; SEC, serous endometrial carcinoma.

### Single-cell census of normal, pre-dysplastic, and dysplastic mouse endometrium in diestrus

Preprocessing steps included principal component analysis and quality control to remove ambient RNA, doublets, and damaged cells (**Figure 1C** and **Supplementary Figure 1**). Once the dataset was fully integrated and main cell types were identified (**Supplementary Figure 2**), we separated and reclustered the samples by SEC stage (normal, pre-dysplastic, and dysplastic) to identify stage-specific cell population dynamics.

Normal (**Figure 2A**), pre-dysplastic (**Figure 2B**), and dysplastic (**Figure 2C**) datasets contained 7,614, 17,212 and 12,717 cells, respectively. Cell types and their states were identified using genes with known cell-type specific expression (**Supplementary Figure 3**).

**Figure 2.**
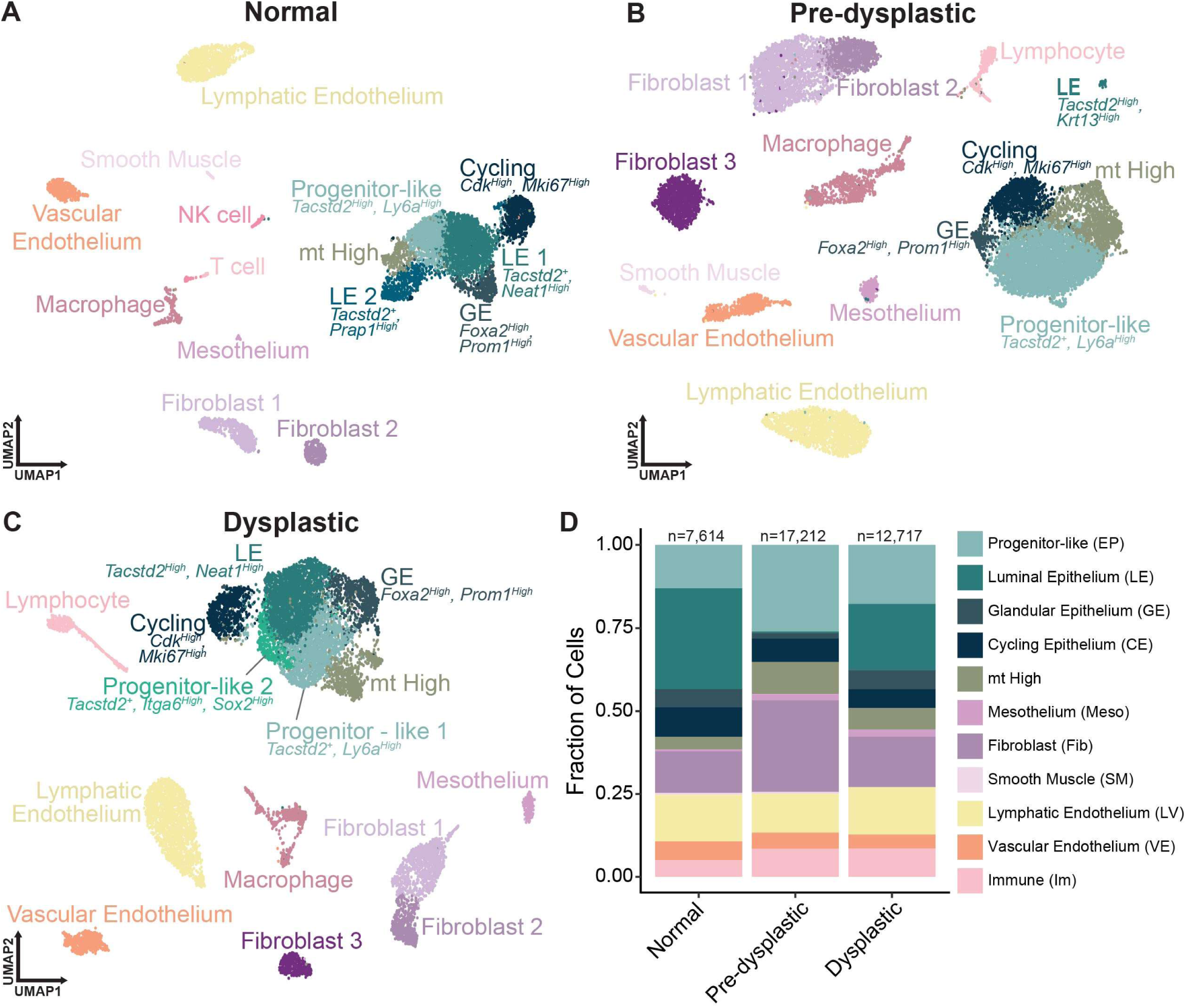
Single-cell census of normal, pre-dysplastic and dysplastic mouse endometrium. (A-C) UMAP visualization of normal (A), pre-dysplastic (B), and dysplastic (C) endometrium. (D) Endometrial cell composition per stage, where cell subtypes were merged to form a single group in the bar chart. mt, mitochondrial; n, number of cells per stage.

The normal dataset included 6 epithelial (*Epcam*+) clusters, including two luminal (*Tacstd2+, Neat1+, Prap1+, Met+*), one glandular (*Foxa2+, Aldh1a1+, Prom1+*), and one cycling (*Mki67+*) cluster. A progenitor-like cluster was also identified, based on expression of stem/progenitor-associated genes, including *Ly6a, Klf4,* and *Sox17*^18–21^. This cluster also expressed *Tacstd2* at high levels, suggesting these cells may be luminal epithelium associated. Additionally, an mt High population was identified, with cells containing a high percent of mitochondrial-associated reads and high expression of other genes related to mitochondrial function (*COX1, COX2*). Further analysis is necessary to determine if this cluster is biologically significant or an artifact of cell preparation. Other cell types identified included fibroblasts (*Dcn+, Col3a1+*), mesothelial cells (*Msln+*), endothelial cells (*Lyve1+, Pecam1+, Vwf+*), smooth muscle cells (*Acta2+*), and immune cells (*Adgre1+*, *Fcgr1/3+*, *Cd3e/4+, Xcl1+*).

Our pre-dysplastic dataset (**Figure 2B**) contained cell types found in the normal dataset. However, only one luminal epithelium cluster was present, while the progenitor-like cluster included many luminal epithelial-associated genes, such as *Tacstd2+* and *Prap1*+, concurrently with *Ly6a, Klf4,* and *Sox17* expression. We have also found the appearance of third fibroblast population (*Dio2+, Rxfp1+,* and *Palld+)* in this stage.

Finally, our dysplastic dataset (**Figure 2C**) showed addition of second progenitor-like clusters marked by *Itga6* and *Sox2* expression. The third, pre-dysplastic fibroblast population persisted in the dysplastic dataset, but additionally showed high expression of *Igf1* and *Des* (**Supplementary Figure 3A**). Instead of previously detected *Acta2+* smooth muscle cell cluster, *Acta2+* cells were present in the fibroblast 1 population at dysplastic stage.

Overall, progression from normal to pre-dysplastic stage was marked by an increase in the frequency of fibroblasts, as well as an increase in the immune cell population, specifically macrophages (**Figure 2D**). In dysplastic stages, the relative size of the fibroblast population returned to near normal conditions. However, the increase in the immune population was maintained.

### Immaturity of luminal epithelium precedes dysplastic changes during carcinogenesis

To further explore the cell-type dynamics within the epithelial compartment, we subsetted and reclustered the epithelial cells of each SEC stage. The normal epithelial subset included 4,689 cells forming seven clusters (**Figure 3A**). Those clusters included two luminal epithelium, one glandular epithelium, one progenitor, one cycling, and one mt High cluster. An additional, putative glandular epithelium cluster was identified based on high expression of *Foxj1* (**Supplementary Figure 3B**), a ciliation marker commonly expressed in the endometrial glands^16^.

**Figure 3.**
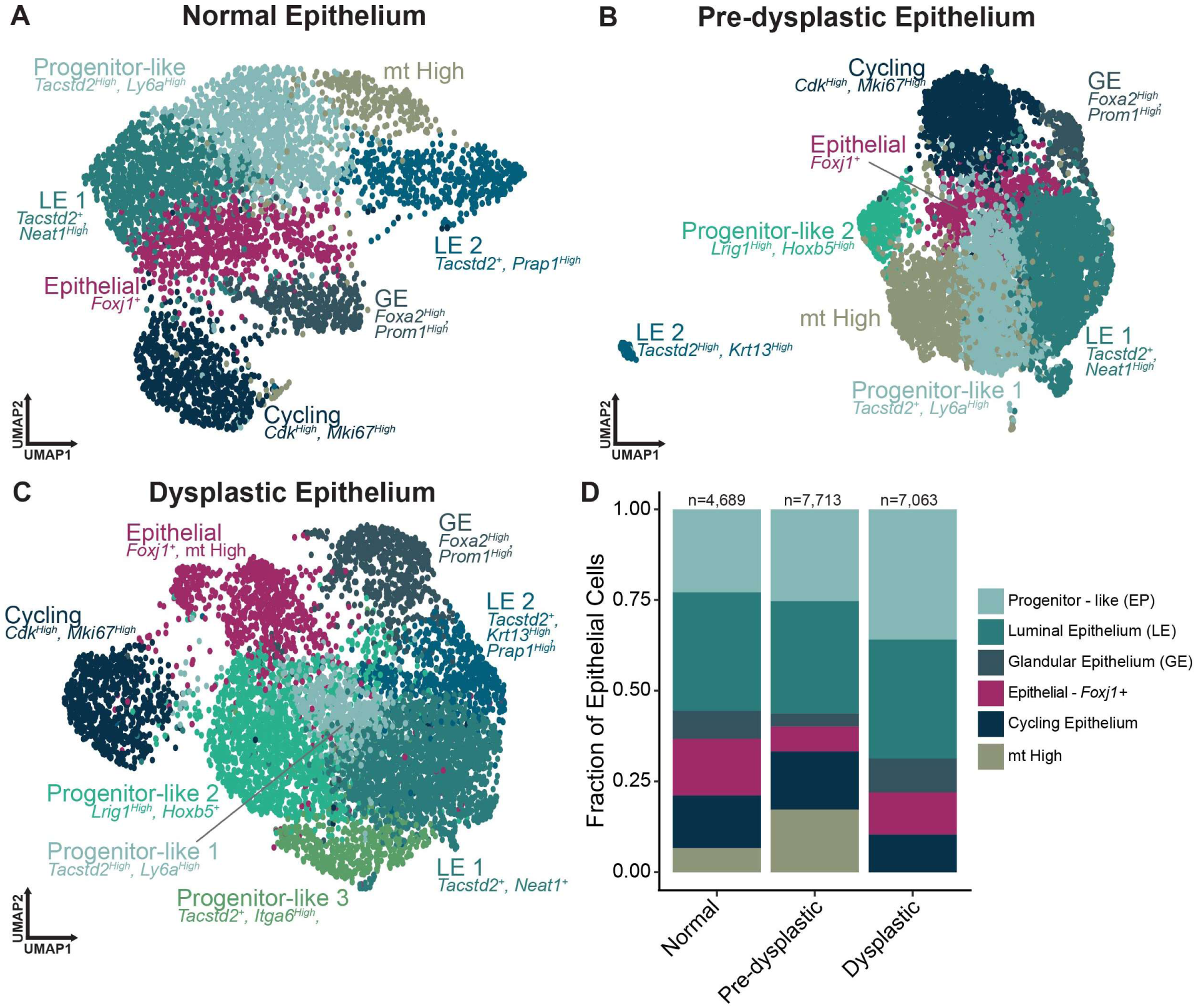
Characterization of normal, pre-dysplastic, and dysplastic mouse epithelial cell states. (A-C) UMAP visualization of normal (A), pre-dysplastic (B) and dysplastic (C) endometrial epithelium. (D) Epithelial cell composition per stage where cell subtypes were merged to form a single group in the bar chart. mt, mitochondrial; n, number of cells per stage.

The pre-dysplastic epithelial subset, comprised of 7,713 cells, contained a diversified progenitor-like populations along with cell types also found in the normal subset (**Figure 3B**). Progenitor-like cells comprised two clusters. One cluster expressed *Tacstd2* along with *Ly6a*, *Klf4, Sox17* and *Tert*^22^ (**Supplementary Figure 3B**). Another cluster also expressed *Tert,* but was marked by high expression of other stemness-associated genes, such as *Hoxb5*^23^ *and Nt5e*^24^, along with *Lrig1,* a marker of quiescence and tumor suppression^25^.

In our dysplastic subset of 7,063 cells, even greater diversification of these progenitor-like populations is observed, with the detection of a third progenitor-like population (**Figure 3C**). This population expressed stem-like genes, *Itga6* and *Sox2*^26^. Furthermore, the previously identified *Foxj1* population has become marked by high expression of mitochondrial-associated genes. Notably, glandular populations always expressed some progenitor-like genes, including *Aldh1a1, Prom1,* and *Lgr5* (**Supplementary Figure 3B**). It is possible that there is a stem-like population within the glandular epithelium cluster that is lowly abundant and unable to be reliably defined at these resolutions.

Consistent with an expectation of a less mature phenotype of transformed cells, there were significant increases in the progenitor population size as SEC progressed (**Figure 3D**). The high frequency of *Tacstd2* positivity in progenitor populations suggests that these immature populations may be associated with the luminal epithelium.

We also compared average expression levels of genes used to identify cell clusters across all SEC stages. Many genes associated with our progenitor-like clusters, such as *Klf4, Sox17, Tert, Nt5e, Hoxb5, Itga6,* and *Sox2,* have higher average expression in dysplastic samples compared to other stages. However, Ly*6a* average expression peaks in pre-dysplastic samples and *Lrig1* expression strongly decreases after SEC induction (**Supplementary Figure 4**). Thus, expression of some progenitor-like genes may not be beneficial for some or all stages of cancer progression.

To further explore pre-dysplastic changes in the endometrium, we evaluated luminal and glandular epithelium in tissue sections (**Figure 4**). Consistent with previous observations^16^, no dysplastic changes were observed until 60 DPI (**Figure 4A**) despite expression of tdTomato in the majority of epithelial cells (**Figure 4B** and **Supplementary Figure 5**). Luminal and glandular epithelial markers were evaluated using TROP2 and FOXA2 co-immunostaining. As reported previously^16^, TROP2 and FOXA2 protein expression was localized to the luminal and glandular epithelium, respectively (**Figure 4C**). However, starting with the pre-dysplastic stage, TROP2 was detected in the glandular epithelium, with some glands co-expressing TROP2 and FOXA2. However, more commonly, TROP2+ glands maintained the typical glandular epithelium cuboidal morphology, while entirely lacking FOXA2 expression. This change to TROP2 expression becomes more pronounced in dysplastic stages (**Figure 4D**), consistent with increase of luminal epithelium immaturity during carcinogenesis.

**Figure 4.**
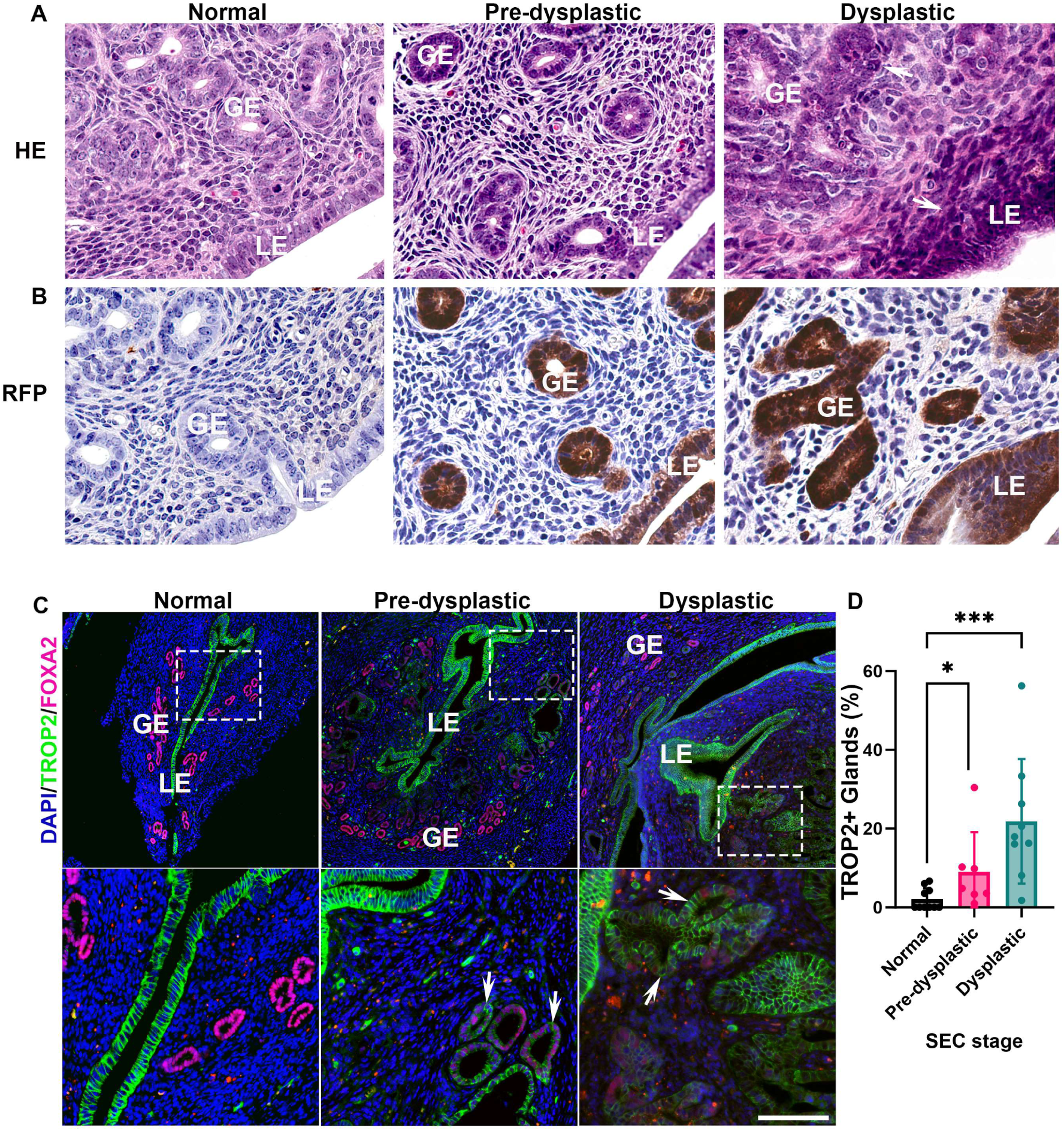
Dysregulation of cell identity during carcinogenesis. (A-C) Cross-sections of the mouse uterus collected at normal, pre-dysplastic, and dysplastic stages. (A) Morphologically normal (normal and pre-dysplastic) and dysplastic (arrows) luminal (LE) and glandular (GE) endometrial epithelium. Hematoxylin and Eosin (HE). (B) Immunohistochemical detection of tdTomato (RFP) expression (brown color) at each stage. ABC Elite method with Hematoxylin counterstaining. (C) Representative images of TROP2 (green) and FOXA2 (pink) expression in the endometrium during SEC formation. Arrows indicate TROP2 expression in glandular structures. Dotted rectangles in top panels denote area magnified and shown in bottom panels. Immunofluorescence with DAPI counterstaining (blue). Confocal microscopy. (D) Quantitative analysis of the frequency of glands with TROP2+ cells compared to total glands in the endometrium in normal (n=10), pre-dysplastic (n=7), and dysplastic (n=9) samples. (A-C) Scale bar in (C) represents 40 μm (A and B) and 250 μm in top panels and 83 μm in bottom panels (C). (D) *P<0.05, ***P<0.001, Mann-Whitney U tests. All error bars denote s.d.

### Cell interaction dysregulation in pre-dysplastic and dysplastic endometrium

To gain additional insights into mechanisms of cell dysregulation during carcinogenesis, we evaluated signaling changes between cells using CellChat^27^. According to this analysis, pre-dysplastic samples were marked by a significant decrease both in the predicted number of interactions and the strength of those interactions (**Figure 5A** and **B**). Interestingly, in dysplastic samples, there was a recovery in the predicted number of interactions, almost back to normal levels. Moreover, the dysplastic samples were more enriched in signaling than normal, as described by the predicted interaction strengths.

**Figure 5.**
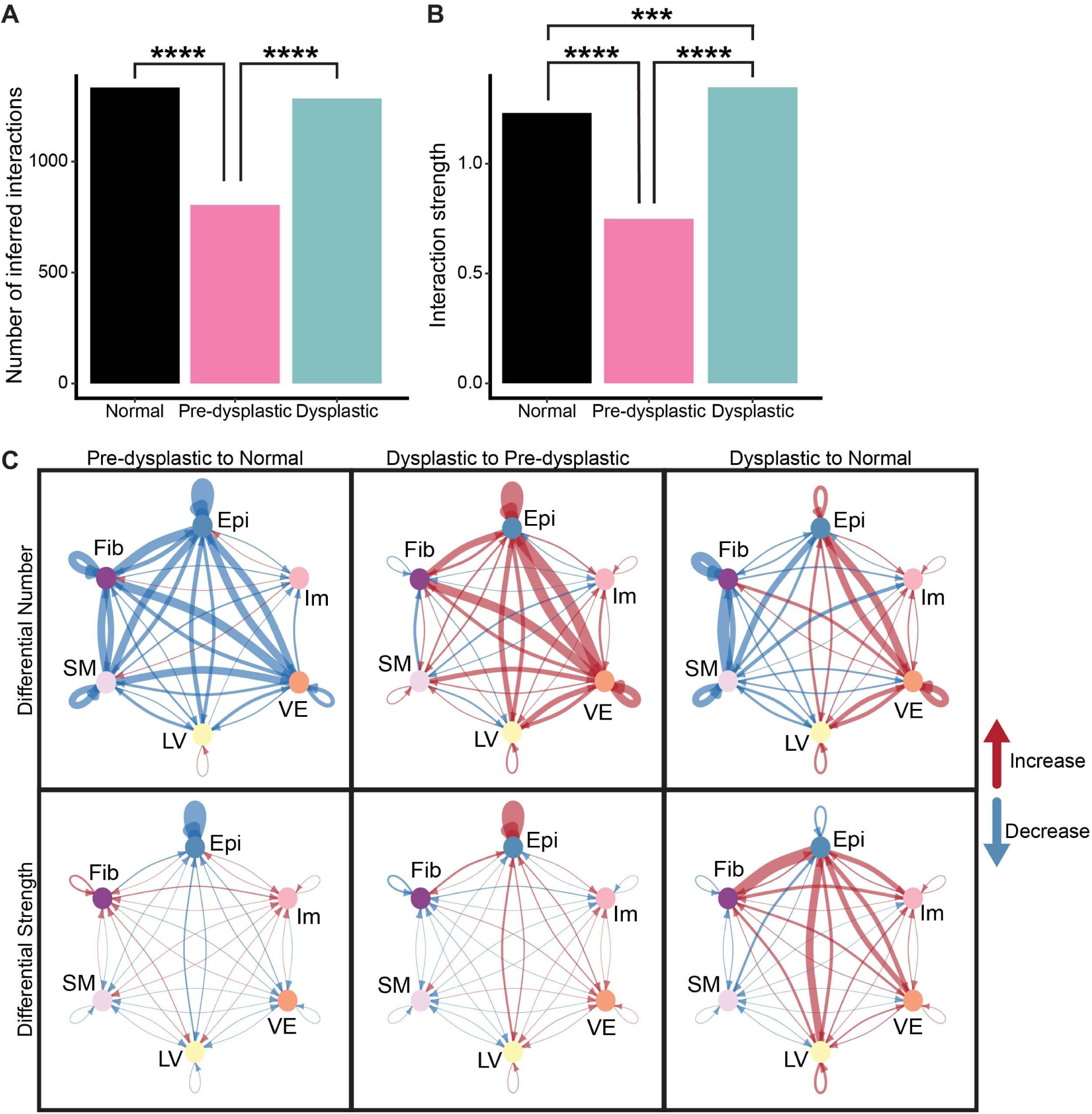
Cell-cell interaction changes during carcinogenesis. (A and B) Quantification of the predicted number (A) and strength (B) of interactions by stage. The strength of interactions is described as the summation of the probabilities of associated ligand-receptor pairs^27^. (C) Circle plots of differential interactions between clusters by stage. Thickness of connecting lines is proportional to the magnitude of differential number/strength of inferred interactions. Red connecting lines indicate an increase in interaction, whereas blue indicates a decrease relative to first listed stage at the top of columns. Epi, Epithelial; Fib, Fibroblast; Im, Immune; LV, Lymphatic Endothelium; SM, Smooth Muscle; VE, Vascular Endothelium. (A-B) ***P<0.001, ****P<0.0001, Fischer’s Exact tests.

To map differential interactions between cell types by stage we have utilized circle plots (**Figure 5C**). As compared to normal samples, the pre-dysplastic stage is marked by a decrease of interactions between all cell types. As compared to pre-dysplastic data, cell-cell interactions in dysplastic stage are significantly increased in their number and strength. This is observed in both inter-epithelial and epithelial-stromal interactions. However, as compared to normal endometrium, dysplastic stage inter-epithelial interactions are only increased in number and not in strength. Furthermore, interactions between epithelial and stromal groups are preferentially increased in endothelial populations.

Taken together, our observations are consistent with observed cellular immaturity during pre-dysplastic stages of carcinogenesis. They also suggest an aberrant character of recovery in cell-cell interactions with progression to the dysplastic stage.

### Comparative screening for early diagnostic and prognostic markers

To identify clinically relevant markers of early-stage disease, we selected 179 genes based on their elevated expression in either pre-dysplastic or dysplastic single-cell RNA sequencing samples compared to normal. Consistent with previous findings, high expression of *Cdkn2a*, a known marker of both mouse and human SEC, was detected^16^ (**Figure 6**). We have also confirmed that expression of *Pgr* was downregulated as reported previously for both mouse and human SEC^4,28^ (**Supplementary Figure 6, Supplementary Figure 7**).

**Figure 6.**
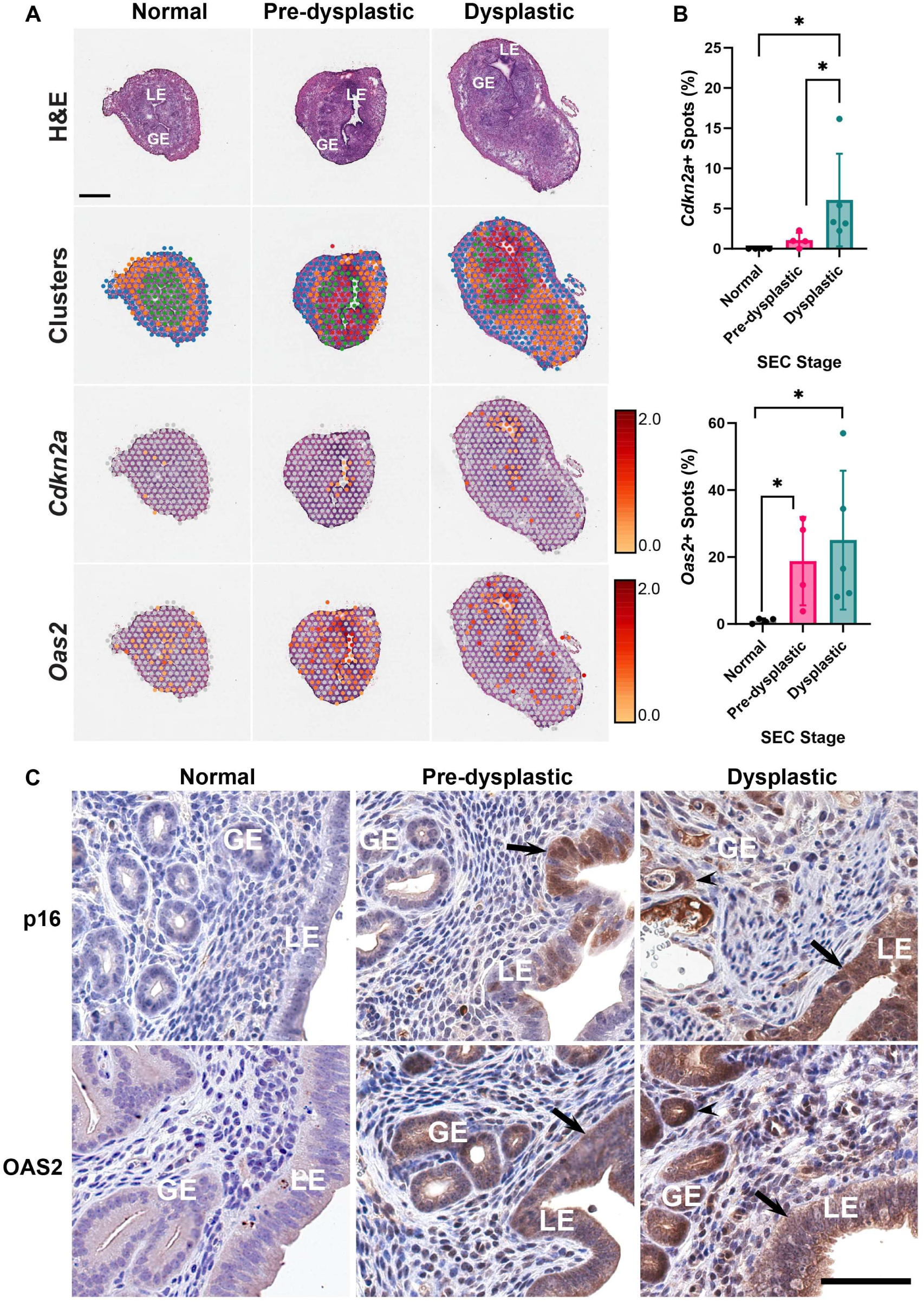
Spatial RNA expression during carcinogenesis. (A) Histological (Hematoxylin and Eosin, H&E) and spatial transcriptomic (Clusters) visualization, and detection of *Cdkn2a* and *Oas2* mRNAs in normal, pre-dysplastic, and dysplastic endometrial epithelium. Visium spatial RNA sequencing. Scale represents the log normalized expression of the spots across each individual sample. (B) Frequency of *Cdkn2a*+ (top) and *Oas2*+ (bottom) spots within the endometrium in normal (n=4), pre-dysplastic (n=4), and dysplastic (n=5) Visium samples. A cutoff of 25% of the maximum log normalized expression value determined spot positivity. *P<0.05, Mann-Whitney U tests. All error bars denote s.d. (C) Immunohistochemical detection (brown color) of p16 (encoded by *Cdkn2a*) and OAS2 (encoded by *Oas2*) expression in glandular epithelium (GE, arrowheads) and luminal epithelium (LE, arrows). ABC Elite method with Hematoxylin counterstaining. Scale bar in (C) represents 615 μm (A) and 60 μm (C).

After screening selected genes for differential expression in our Visium specimens, we identified 45 genes differentially expressed between normal and SEC samples. We tested human homologs of these genes for association of their expression with overall and relapse-free survival of endometrial carcinoma patients by using Human Protein Atlas and KM Plotter^29^ (**Supplementary Table 1**). Expression of *CDKN2A* was used as a diagnostic marker typically associated with SEC (**Figure 7** and **Supplementary Figure 8**). The concordant expression of the identified genes with *CDKN2A* had strong association with worst prognosis. Based on these criteria, expression of 44 out of 45 genes correlated with worst patient survival. According to highly reproducible pattern of their expression across individual samples in our mouse model, four genes were identified as promising targets for further comparative evaluation of their functions in mouse/human serous endometrial carcinogenesis. Those included already known SEC markers, *CDKN2A* and *PGR*, and two novel markers associated with innate immunity^30^, *OAS2* and *OASL* (mouse *Oasl2*; **Figure 6** and **Supplementary Figure 7**).

**Figure 7.**
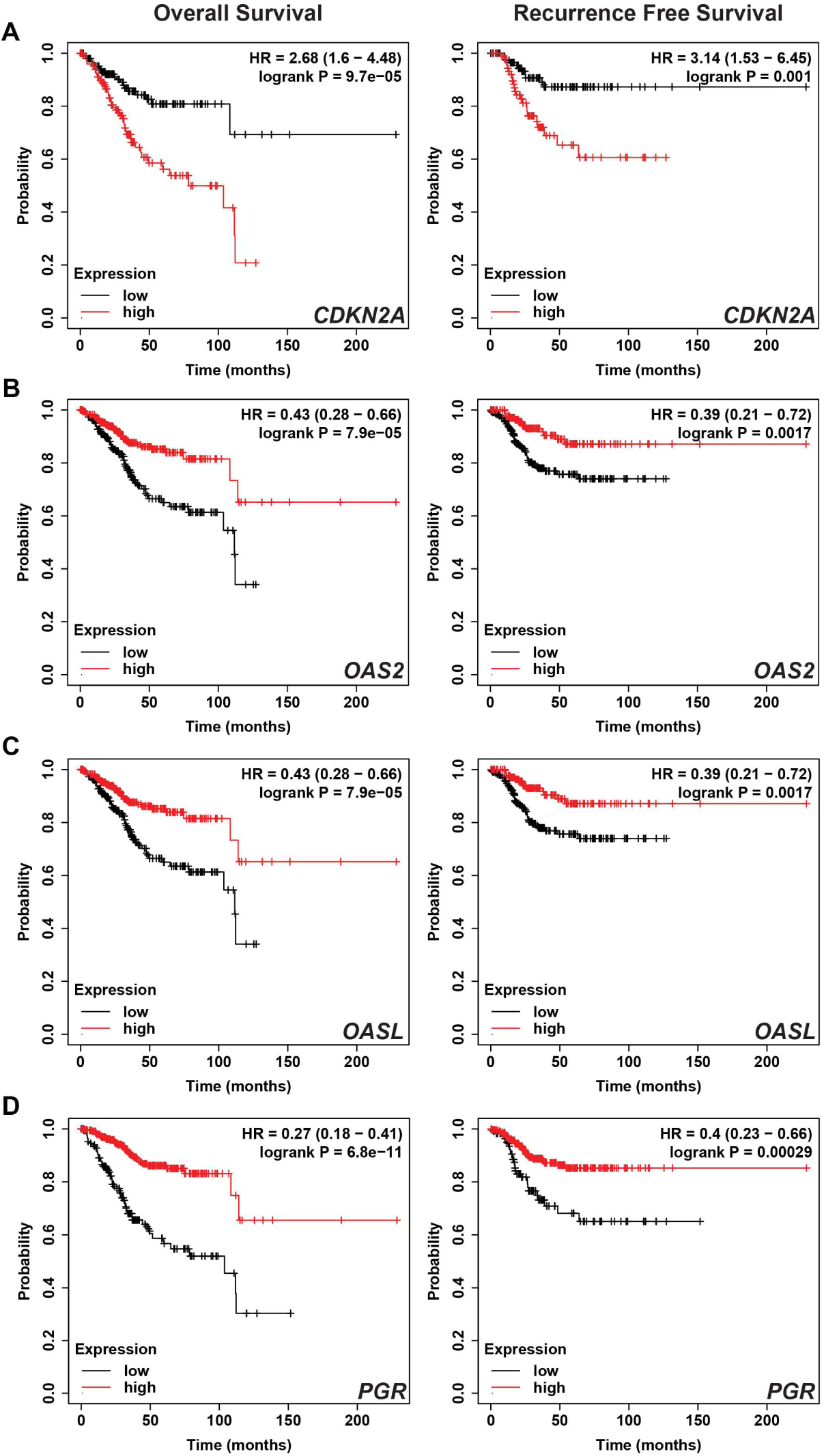
Survival curves for diagnostic genes. (A-D) Kaplan-Meier plots for mRNA expression of *CDKN2A* (A), *OAS2* (B), OASL (C) and PGR (D) with overall survival (left) and recurrence free survival (right). (A) *CDKN2A* survival was determined by a cohort of grade 3 endometrial carcinoma samples. (B-D) Survival was stratified using a ratio of the gene of interest to *CDKN2A*, using the latter genes as a diagnostic marker of SEC. Concordant expression of both genes are expressed highly leads to ratio one, falling in the low ratio category. Data collected from KMPlotter^29^.

## Discussion

We used a mouse model of SEC previously established in our lab to create a single-cell census of serous endometrial carcinogenesis. To identify cell dynamics related to carcinogenesis, we utilized three timepoints in carcinogenesis: normal, pre-dysplastic, and dysplastic. According to our results, the luminal epithelium shows immature properties already in pre-dysplastic stage, before morphologically atypical cells can be detected. With progression to dysplastic stage, further increase in proportion and diversity of immature progenitor-like population expressing luminal markers is observed. However, the original progenitor-like population decreases in abundance and is overtaken by the two other SEC-associated, progenitor-like populations. Furthermore, diversity of such immature cells also increases, in agreement with expected increase in heterogeneity of overt malignancies. Notably, a new population of cells with stemness markers (*Itga6+, Sox2+*) is observed in dysplastic samples. It will be interesting to test if such cells represent nascent cancer propagating cells, also known as cancer stem cells.

Consistent with observed transcriptomic changes indicative of emerging immature phenotype, we detected an increased expression of the luminal and progenitor epithelial cell marker, TROP2, in the glandular epithelium already at pre-dysplastic stage. Taken together, our studies suggest an important role of the luminal epithelium in SEC pathogenesis. Previously it has been reported that conditional activation of *Wnt* signaling in conjunction with expression of the oncogenic form of *Pik3ca* in *Axin2-*expressing cells of the glandular epithelium leads to formation of endometrial carcinoma^31^. Since alterations in Wnt and Pik3ca signaling are commonly associated with endometrioid endometrial carcinoma, our work offers an opportunity to test if serous and endometrioid endometrial carcinomas have different cells of origin. Further cell lineage fate-based studies should explore this possibility.

Our CellChat analysis showed a significant decrease of cell-cell interactions during pre-dysplastic stage. This observation suggests that an association between dysregulated cell-cell connection and immature cell status is an early step in serous endometrial carcinogenesis. Interestingly, cell-cell interactions again increase with cancer progression, albeit with an overall different pattern and strength of interactions. Preferential increases of interactions between epithelial and stromal cells, especially endothelial populations, open new opportunities to identify clinically relevant ligand-receptor interactions during serous endometrial carcinogenesis. This may offer a possibility to explore pre-dysplastic cell-signaling vulnerabilities for the development of preventive medicine approaches.

By using a multi-level approach combining single-cell and spatial transcriptomics paired with screening for clinically relevant genes in human endometrial carcinoma, we were able to identify a panel of 44 genes suitable for further testing of their validity as early diagnostic and prognostic markers. As an example of this approach, we have identified two novel SEC-associated genes, *Oas2* and *Oasl2* (*OAS2* and *OASL* in humans), with *OAS2* being a most promising marker for early SEC detection. *Oas2* and *Oasl2* are members of 2-5A synthetase family of enzymes that are part of the innate immune response to viral infections^30^ and have been linked to immune cell infiltration in cancer. Association of OAS genes with cancer outcomes are tissue-type specific. For example, high expression of *OAS* family genes in pancreatic adenocarcinoma^32^ is predictive of poor patient prognosis, however *OAS2* expression is associated with favorable outcomes in breast cancer^33^ and colorectal cancer^34^.

SEC tumors frequently have low expression of immune markers, including those often used to predict immunotherapy response (microsatellite instability and PDL1)^5,35^. It has been reported that upregulation of *OAS2* can cause defects in T cell signaling by downregulating CD3 expression in T cells in other diseases^36,37^. A few studies investigating immune response in endometrial carcinoma have been done but are infrequently stratified by histological subtype, leading to conflicting results. Further studies of OAS genes in our model may facilitate resolving this problem.

In summary, our findings provide a genome-wide overview of single-cell and spatial RNA transcriptomic changes during early stages of SEC. These studies show that anticipated decrease of differentiated cell states is associated with a loss of cell-cell interactions. Such loss is transient and is followed by formation of a new set of neoplastic features expected to facilitate further cancer progression. Our study also shows feasibility of using a genetically defined mouse model to identify novel prognostically significant genes in human SEC.

## Methods

### Experimental animals

Tg(Pax8-rtTA2S*M2)1Koes/J (Pax8-rtTA) Tg(tetO-Cre)1Jaw/J(Tre-Cre) mice (Perets et al., 2013) mice and *Gt(ROSA)26Sor^tm9(CAG-tdTomato)Hze^* (Ai9) mice (JAX stock no. 007909) were obtained from The Jackson Laboratory (Bar Harbor, ME, USA). *Trp53^loxP/loxP^* and *Rb1^loxP/loxP^* mice, which have *loxP* alleles flanking their respective genes, *Trp53* and *Rb1,* were a gift from Dr. Anton Berns (The Netherlands Cancer Institute, Amsterdam, The Netherlands). For all experiments mice were collected in later diestrus, also described by some as early proestrus^38–41^. Briefly, criteria for this stage included observing almost exclusively leukocytes in vaginal smears, a medium-wide lumen, dense/early edematous stroma, and medium-high proliferation of the glandular and luminal epithelia (Ki67 index 10-95%). All experiments and maintenance of the mice were performed following ethical regulations for animal testing and research. The Cornell University Institutional Animal Care and Use Committee (IACUC) approved all animal protocols, and experiments were performed in compliance with its institutional guidelines.

### Doxycycline induction

Doxycycline was administered through a single intraperitoneal injection (i.p.) to Pax8-rtTA Tre-Cre *Trp53^loxP/loxP^Rb1^loxP/loxP^* Ai9 mice and control mice at 6 weeks to 6 months of age. Doxycycline was administered at a dose of 12 μl g^-1^ body weight at a concentration of 6.7 mg ml^-1^ in sterile PBS. Mice were identified to be in diestrus phase of the estrous cycle before induction using vaginal smears. All mice were euthanized by CO_2,_ and further analyses were carried out.

### Pathological evaluation

All mice underwent gross pathology evaluation at the time of necropsy. Potential sites of endometrial carcinoma metastasis were evaluated carefully, including the local lymph nodes, omentum, peritoneum, and lungs.

### Histology, immunohistochemistry, and image analysis

All tissues were fixed in buffered 4% paraformaldehyde overnight at 4 °C followed by standard tissue processing and paraffin embedding. Histology and immunohistochemistry stainings were carried out on 4-μm-thick tissue sections. For immunohistochemistry, antigen retrieval was performed as needed by incubation of deparaffinized and rehydrated tissue sections in boiling 10 mM sodium citrate buffer (pH 6.0) for 15 min. All primary antibodies used for immunostaining are listed in Supplementary Table 2. The primary antibodies were incubated at 4°C overnight or 1 hr at room temperature, followed by incubation with secondary biotinylated or fluorophore conjugated antibodies (one hour, at room temperature, RT). Incubation with biotinylated antibodies slides followed by modified Elite avidin-biotin peroxidase (ABC) method (Vector Laboratories, Burlingame, CA, USA; pk-6100) for 30 minutes (RT). Stained sections were scanned by ScanScope CS2 (Leica Biosystems, Vista, CA) with a 40X objective, followed by the analysis with the ImageJ software (National Institutes of Health, Bethesda, MD, USA). Following Vectashield Vibrance Antifade Mounting Medium with DAPI (H-1800, Vector Laboratories, Newark, CA), immunofluorescence samples were tilescanned (Zeiss 710 upright Confocal, Cornell University’s Biotechnology Resource Center) at 40X. Analysis was performed with ImageJ software.

### Single-cell isolation

For collection, induced mice were sacrificed at various time points in the diestrus phase of the estrous cycle. Each mouse was collected and processed independently to generate single-cell suspensions. Both uterine horns of individual mice were collected and placed in sterile 1X PBS containing 100 IU ml^-1^ of penicillin and 100 μl ml^-1^ streptomycin (Corning, 30-002-Cl). A portion of one uterine horn per mouse was set aside in 4% paraformaldehyde at 4°C for paraffin embedding. The remaining uterine horns were placed in a 200 μL drop of the same PBS solution, cut open lengthwise, and minced into 1.5-2.5 mm pieces with scalpels. Minced tissues were transferred with the help of a sterile, wide bore 200 μl pipette tip into a 15 ml centrifuge tube containing 2 ml of the same PBS solution and then centrifuged for 6 minutes at 400 rcf at 4°C. Then, the minced tissues for individual mice were digested using a Papain Dissociation System (Worthington Biochemical Corporation, New Jersey). Tissue was digested in papain mixture from the kit at 37°C for 1.5 hours with periodic mechanical perturbation. After papain digestion, the papain was inhibited with 3 ml of our stop solution, a DMEM/FBS solution (DMEM Ham’s F12, Corning 10-092-CV; 20% fetal bovine serum [FBS], Sigma-Aldrich F4135; and 0.1 mg ml^-1^ DNase I, Stem Cell Technologies 07900). Cell suspension was placed in a new tube. Cells were centrifuged as before, and the supernatant was removed. Next, 1.35 ml of the albumin-ovomucoid inhibitor provided by the kit was added to the suspended cells, and the cells were resuspended gently. The reaction was stopped using 3 ml of our stop solution, mixed, and then cells were spun down at the same rate as before. The supernatant was removed, and the pellet was resuspended in 2 ml of 0.25% TrypLE Express Enzyme solution (Invitrogen, 12604013) prewarmed to 37°C. The tube was then incubated at 37°C with a loose cap for 10 minutes in a 5% CO_2_ incubator. Cells were resuspended with a 1-ml pipette tip 40 times, and then digestion was inhibited with our stop solution. Cells were spun down and supernatant was removed before the addition of 1 ml Dispase-mUE (Dispase II [7 mg ml^-1^ DMEM Ham’s F12], Neutral Protease, Worthington NPRO2; and 10 μg μL^-1^ DNase). Cells were aspirated in the solution gently 40 times using a 1 ml pipette tip. Then suspensions were filtered through a 35 μm filter (Falcon 08-771-23) to remove debris, and remaining cells were spun down. Cells were resuspended in 100 μl A-scR-Medium (5% FBS in DMEM Ham’s F12 with 1 mM Y-27632 [Toris 1257], placed on ice and transferred to Cornell’s Genomics Core for single-cell RNA sequencing for library preparation.

### Single-cell RNA sequencing library preparation

To begin single-cell RNA sequencing library preparation, cell aliquots were stained with trypan blue for live and dead cell calculation. Live cell preparations with a target cell recovery of 5,000 cells were loaded on a Chromium Controller (10X Genomics, Single Cell 3’ v3 chemistry) to perform single-cell partitioning and barcoding using the microfluidic platform device. The NextSeq500 System was used to sequence the cDNA library samples after barcode preparation.

### Download and alignment of RNA sequencing data

For sample sequence alignment, a custom reference for mm39 which included the Ai9-*tdTomato* gene annotation was built using the CellRanger (v7.1.0, 10X Genomics) *mkref* function. From UCSC Genome Browser downloads, mm39.fa soft-masked assembly sequence and mm39.ncbi.RefSeq.gtf file for gene annotations (last modified 2022-11-30) were downloaded for the reference. These files were appended with the Ai9-TdTomato annotation. The Ai9-tdTomato sequence included both the tdTomato sequence and WPRE sequence and was annotated as TdTomato-UTR. The fasta and gtf files for these sequences were shared with us from the Baker Lab at the University of Edinburgh^42^. These files were concatenated with the mm39 fasta and gtf files. Finally, raw reads were aligned to the mm39+tdTomato reference genome using CellRanger (v7.0.1, 10X Genomics).

### Single-cell RNA sequencing preprocessing and batch correction

First, SoupX (v1.6.2; github.com/constantAmateur/SoupX) was used to remove ambient RNA signals using their default workflow (*autoEstCounts* and *adjustCounts*). The standard Seurat (v4.3.0) workflow (*NormalizeData, ScaleData, FindVariableFeatures, RunPCA, FindNeighbors, FindClusters, and runUMAP*; github.com/satijalab/seurat) was used for Seurat object preparation and preprocessing. DoubletFinder (v2.0.3; github.com/chris-mcginnis-ucsf/DoubletFinder) was used to predict doublets in each dataset individually. The default *P_N_* value (0.25) was used, and BC_mvn_ optimization was used to identify ideal *P_K_* values for each sample. Doublet rate was informed by the 10x Chromium Handbook. Then, all datasets were merged. Cells with fewer than 200 features (nFeature), fewer than 750 transcripts (nCounts), or more than 25% of unique transcripts related to mitochondrial genes, and predicted doublets were removed. After preprocessing and merging of the datasets, batch correction was performed using Harmony (v0.1.1; github.com/immunogenomics/harmony). We then used Seurat to process the integrated data.

### Clustering parameters and annotations

Cell types were determined using multiple subsets: as a fully integrated object and as individually integrated objects based on SEC stage (normal, pre-dysplastic, and dysplastic). For each combination of samples, principle component analysis was completed and the dimensions accounting for 95% of variability in the datasets were used to generate SNN graphs (*Seurat::FindNeighbors*). Cell clustering was determined using Louvain clustering on the output graphs (*Seurat::FindClusters*). Clustering resolution and K values were adjusted for each combination of samples to best represent the data. Cell types were determined based on canonical genes. If needed, clusters that had similar canonical gene expression patterns underwent differential expression analysis (*Seurat::FindMarkers*) for better identification. Cell types with multiple associated clusters were also merged to show combined expression for some plots.

### Differential gene expression analysis

For differential gene expression analysis, the fully integrated dataset including all samples was used. The dataset was subsetted (*Seurat::subset*) to reflect the cell type of interest for each particular plot. Epithelial subsets represent the Luminal Epithelium, Glandular Epithelium, Epithelial Progenitors, and cycling populations. *Seurat::FindMarkers()* was used to identify the top 100 differentially expressed genes between the groups of interest.

### Cell-cell interactions analysis with CellChat

Cell interaction analysis and visualization were performed using CellChat (v1.6.1, github.com/sqjin/CellChat). The cell-type labels from initial Harmony integration from the fully integrated single-cell RNA sequencing dataset were used. To simplify identification of interaction patterns, similar cell types were combined into a single cluster. For example, all epithelial cells were combined into Epi cluster. The mesothelium population was removed, as it is unlikely that the cells would have large interactions with endometrial cell types. Default parameters were used when necessary.

### Visium spatial RNA sequencing sample preparation

Mouse uteri from Doxycyline induced Pax8-rtTA Tre-Cre *Trp53^loxP/loxP^Rb1^loxP/loxP^* Ai9 experimental mice and control littermates were dissected and frozen in O.C.T. (Tissue Tek O.C.T. Compound, VWR, 25608-930) filled molds (Disposable Based Molds, 15 x 15 mm, VWR, 60872-488) on a metal block chilled in dry ice for 10 minutes. Molds were wrapped in aluminum foil, placed and sealed in zip lock plastic bags and stored at -80°C (10xGenomics, protocol CG000240RevD). Embedded samples, and all materials for cryosectioning were equilibrated in a cryostat (Leica CM1950) chamber temperature - 21°C (object head, -16°C) for 30 minutes. Transverse uterus sections of 10µm thickness were placed on chilled Visium Tissue Optimization Slides (10xGenomics, 1000191 Slide Kit), and Visium Spatial Gene Expression Slides (10xGenomics, 1000187 Slide Kit). Tissues processed on the Visium Tissue Optimization Slides (10xGenomics, protocol CG000238RevD) indicated the optimal uterine tissue permeabilization time was 12 minutes. For tissue optimization experiments Tiff fluorescent images (Microscope Olympus EX51, Camera Olympus XM10) were captured with a TRITC filter cube, 2x objective, and 700-ms exposure time. Uterine tissues on Visium Spatial Gene Expression Slides were processed for Hematoxylin & Eosin (H&E) staining (10xGenomics, protocol CG000160RevC). Immediately after staining, digital brightfield histology images were taken at 40x (Leica BioSystems, Aperio ScanScope CS2). Without delay cDNA Synthesis, Second Strand Synthesis & Denaturation, and cDNA Amplification was performed (10xGenomics, protocol CG000239RevE). Samples were frozen at -20°C and transferred to Cornell Genomics Facility for Quality Control (QC), Gene Expression Library Construction and sequencing.

The Illumina Sequencing parameters for all Visium samples were as follows, visium library preparations were run 2 times, they were re-balanced and re-pooled in between the 2 runs. The estimated dPCR was 2.5 nMol. Samples were run on a NextSeq2K P2 100bp flowell instrument, sequencing Kit NextSeq 2K P2 100 bp, read length 28+10+10+90, sequencing primer type TruSeq Compatible DNA/RNA, barcode type Dual barcode i7 and i5.

### Visium Spatial RNA sequencing Raw Data Processing

Raw FASTQ files and histology images (Brightfield,16-bit Jpeg, 2025 x 2074 pixels, 16 µm, hourglass dots of fiducial frame in upper left corner) were processed by SpaceRanger (v2.1.1, 10X Genomics). For samples S01 through S12, LoupeBrowser (V6.2.0, 10X Genomics) was used to manually select relevant barcodes for samples that were mounted to an acquisition area, and the provided JSON file of barcodes utilized with the *loupe-alignment* argument in the SpaceRanger counts function. The raw data from our Visium samples were aligned to our custom reference genome of mm39+TdTomato, the same as our single-cell RNA sequencing samples.

### Spatial RNA sequencing preprocessing

Aligned files were opened into LoupeBrowser (V7.0.1, 10X Genomics), where clustering occurred. Low quality spots were identified using a UMI count cutoff of 1000 and Feature count cutoff of 500, these same spots were removed, and the samples were reclustered. All further analysis was performed in LoupeBrowser with log normalized expression values for each individual sample.

### Visium Spot Quantification

After genes of interest were identified, the maximum log normalized expression values for each individual gene across all samples was identified. A 25% of maximum expression cutoff set to minimize variance across the majority of selected genes, as well as show a balanced view of changes in number of positive spots with changes in expression level per spot.

To quantify clusters that were associated with the endometrium according to the H&E staining of individual samples, all barcodes except those in the endometrial clusters were removed. Any spot that remained positive for each individual sample after the expression cutoff was applied was counted. This value was compared to the total number of spots within the endometrium to identify a percent of positive spots for each gene.

### Statistical analysis

Statistical comparisons were performed using a Mann-Whitney U test with GraphPad InStat 3 and GraphPad Prism 10 software (GraphPad Software Inc., La Jolla, CA, USA).

## Data Availability

The single-cell RNA-seq and Visium RNA-seq data reported in this paper is deposited in Gene Expression Omnibus (GEO); accession number (in process). The Source Data provides data for all results requiring quantification. Any additional data supporting the findings of this study are available from the corresponding author upon reasonable request.

## Code Availability

The Seurat objects for each individual SEC stage and Visium RNA-seq cloupe files will be available via download on Dryad upon publication. Epithelial subsets of the respective datasets will also be available through the same platform. All codes for data preprocessing and figure generation will be made available through GitHub upon publication.

## Acknowledgements

We thank Peter A. Schweitzer, Director of the Cornell Genomics Facility, and Ann Tate, Project Manager of Transcriptional Regulation and Expression facility for their invaluable assistance with single-cell RNA sequencing and Visium transcriptomics, and Md Mozammal Hossain, Evelyn Kim, and Eric Lim for their excellent technical support. We also thank David McKellar for his code availability and assistance with learning methods for sequencing analysis in R. This work has been supported by NIH grants (CA248524 and CA260115) to AYN and Cornell Vertebrate Genomics seed funding to AF-N.

## Author contributions

AFN, MP, BDC and AYN designed experiments. AFN, MP, and CSA performed experiments, MP, and BDC carried out bioinformatics analyses, LHE and AYN performed pathological evaluations, AFN, MP, and AYN wrote the paper.

