## Supplementary Information for "Dysregulation of cell state dynamics during early stages of serous endometrial carcinogenesis"

Flesken-Nikitin et al.

### Supplementary Figures

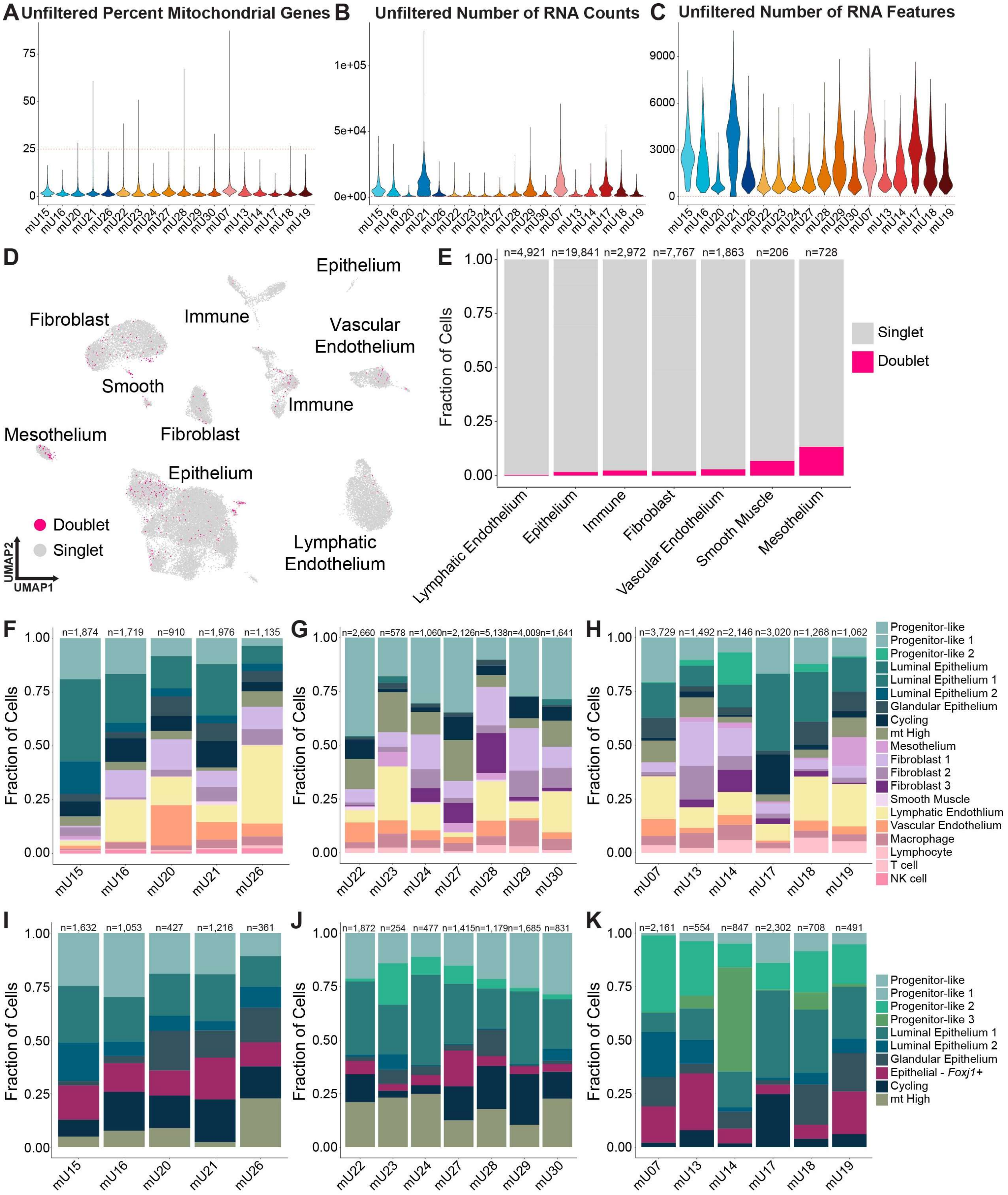


**Supplementary Figure 1. Quality control of single-cell RNA sequencing datasets.** (A-C) Violin plots displaying the unfiltered values for percent mitochondrial genes (A), RNA counts (B), and RNA features/UMI (C) for the cells of each individual sample. A red dotted line represents the filtering cutoff for each respective feature. mU, mouse uterus sample number. (D) A UMAP displaying the clusters in which cell doublets were detected. (E) The percentage of cells in each cluster that were identified as doublets. n, number of cells per cluster. (F-H) The cell type composition of each replicate in the full datasets of normal (F), pre-dysplastic (G), and dysplastic (H) samples. (I-K) The cell type composition of each replicate in the epithelial subsets of normal (F), pre-dysplastic (G), and dysplastic (H) samples. n, number of cells per sample.

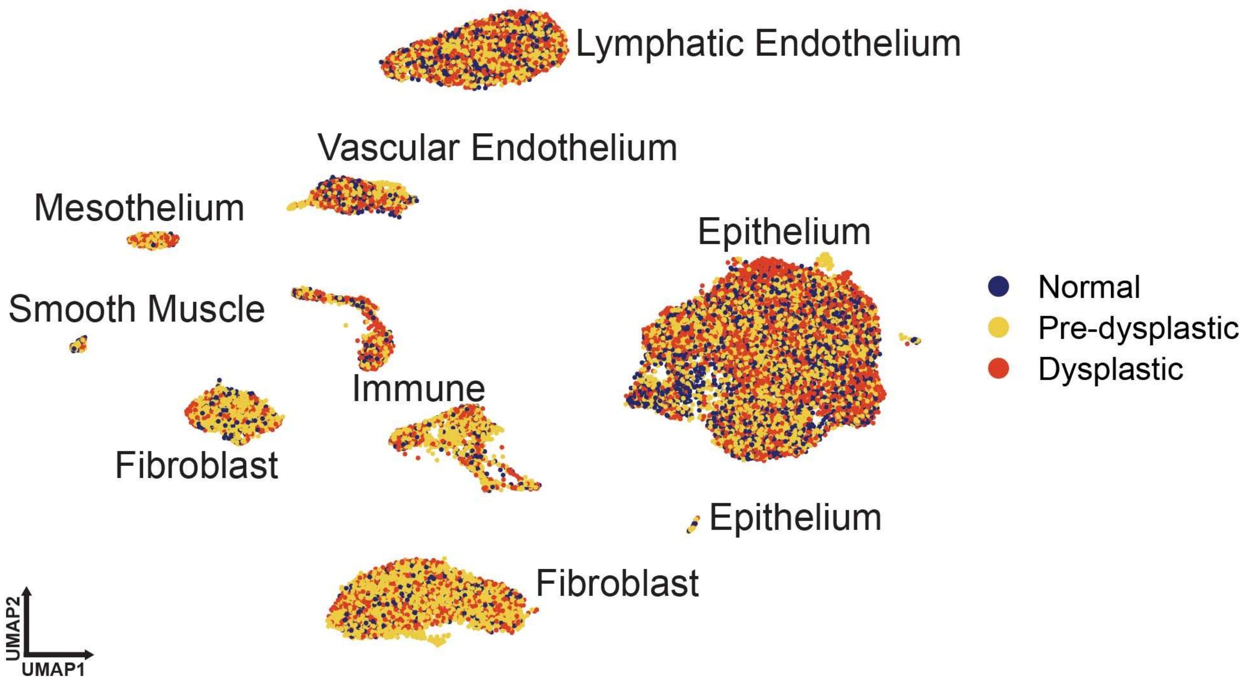


**Supplementary Figure 2. Harmony integrated data set of normal (blue), pre-dysplastic (yellow) and dysplastic (red) samples.**

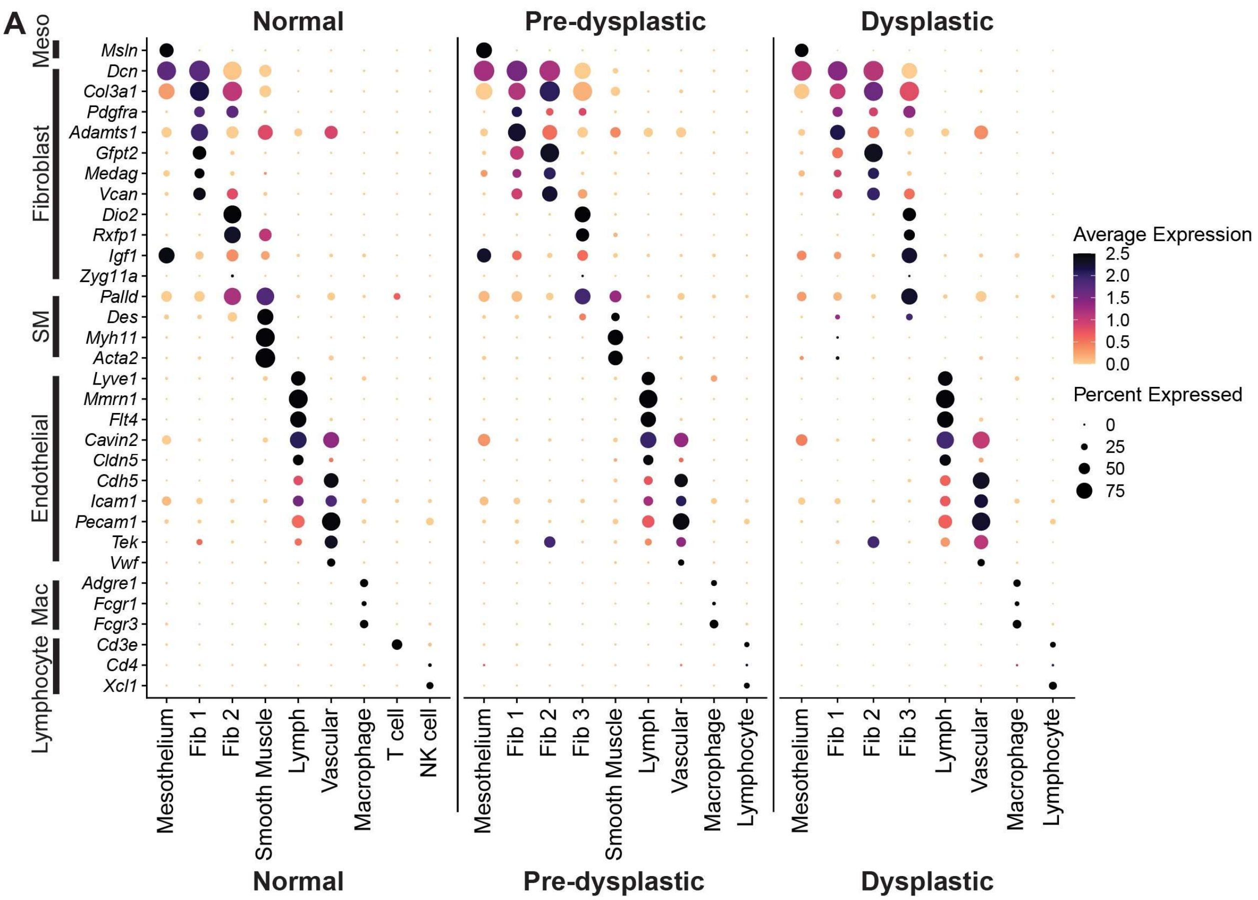
A
Normal
Pre-dysplastic
Dysplastic
Meso
Fibroblast
SM
Endothelial
Mac
Lymphocyte
Msln
Dcn
Col3a1
Pdgfra
Adamts1
Gfpt2
Medag
Vcan
Dio2
Rxfp1
Igf1
Zyg11a
Palld
Des
Myh11
Acta2
Lyve1
Mmrn1
Flt4
Cavin2
Cldn5
Cdh5
Icam1
Pecam1
Tek
Vwf
Adgre1
Fcgr1
Fcgr3
Cd3e
Cd4
Xcl1
Mesothelium
Fib 1
Fib 2
Fib 3
Smooth Muscle
Lymph
Vascular
Macrophage
T cell
NK cell
Lymphocyte
Average Expression
2.5
2.0
1.5
1.0
0.5
0.0
Percent Expressed
0
25
50
75

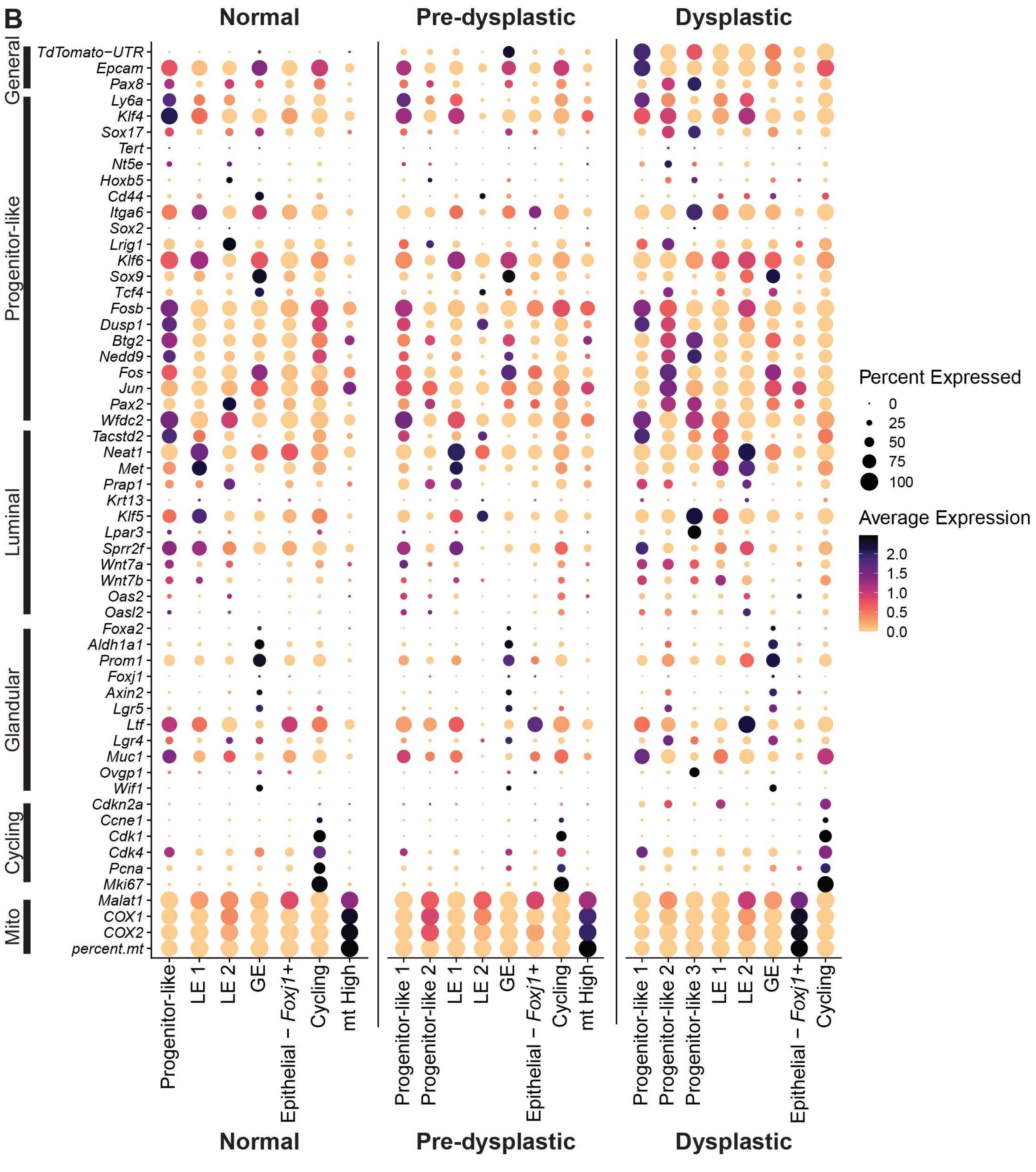


**Supplementary Figure 3. Genes used for cluster naming.** (A) Dot plots showing expression of the genes used to name stromal clusters within each SEC stage. (B) Dot plots showing expression of genes used to name clusters within the epithelial subsets of each SEC stage. Fib, Fibroblast; Mac, Macrophage; Meso, Mesothelium; Mito, Mitochondrial; SM, Smooth Muscle.

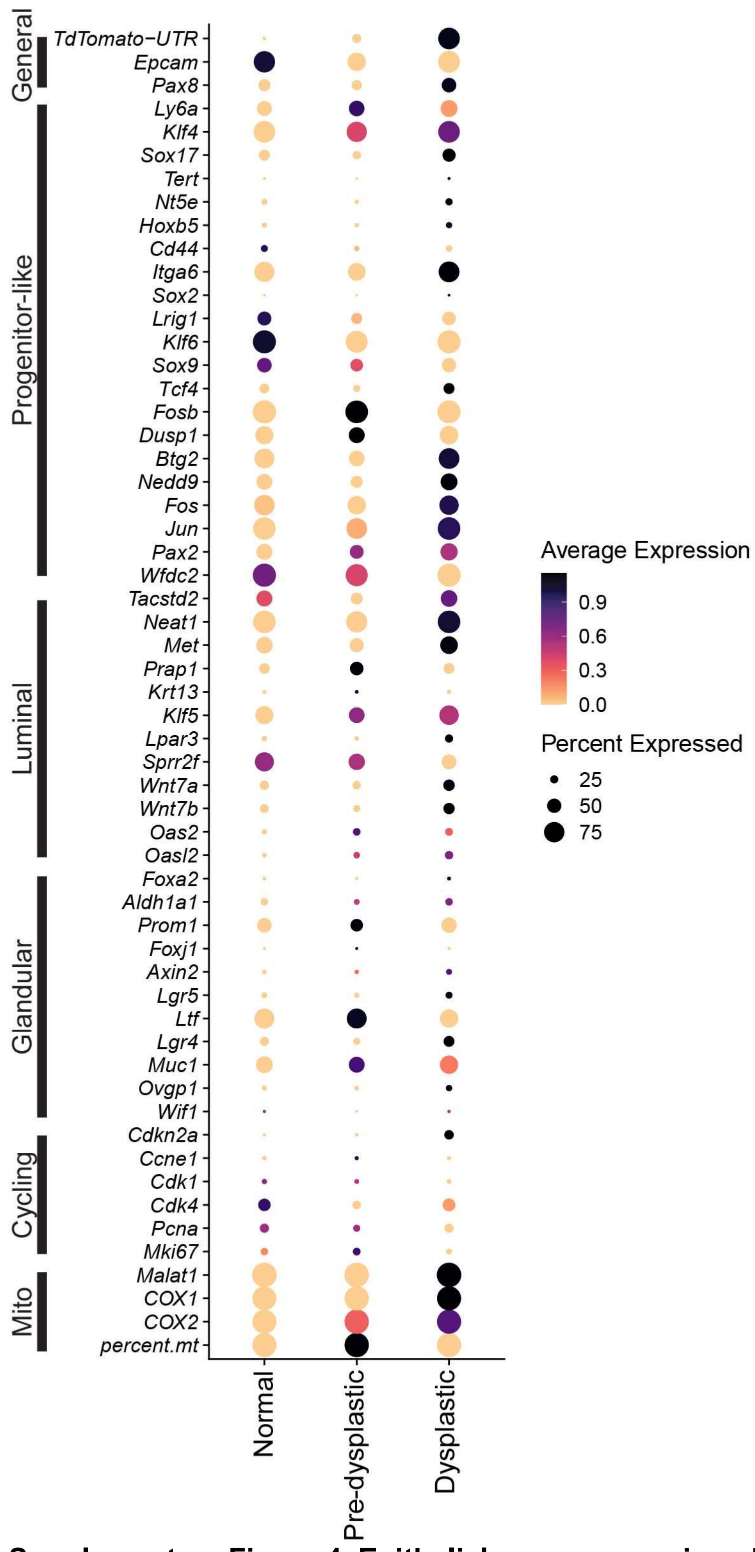


**Supplementary Figure 4. Epithelial gene expression changes during carcinogenesis.** A dot plot showing the comparison of average expression of genes used to name the epithelial clusters across the three stages of SEC.

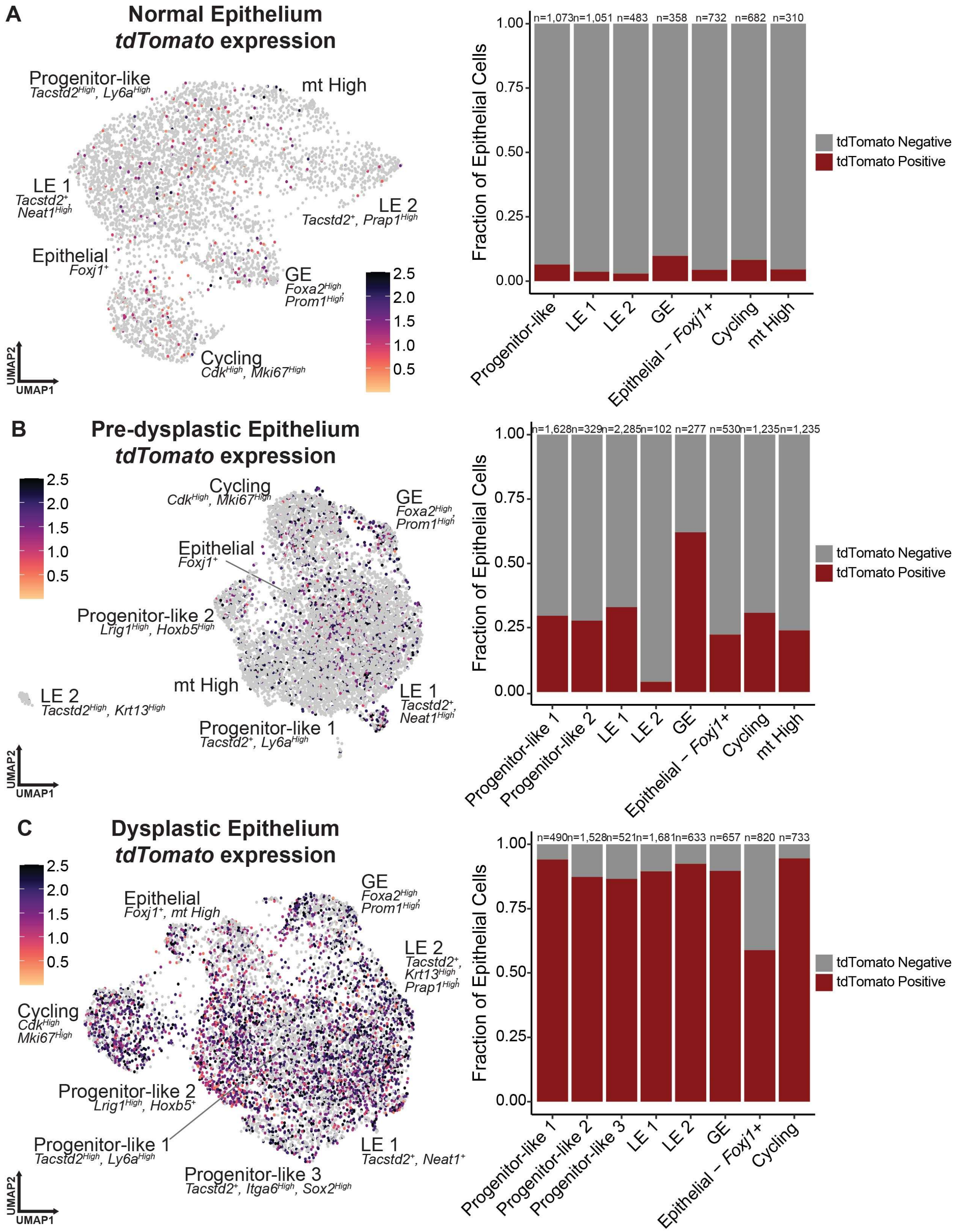


**Supplementary Figure 5. tdTomato expression during SEC formation.** (A-C) Feature plots and bar charts showing *tdTomato* expression in the epithelial subsets of normal (A), pre-dysplastic (B), and dysplastic (C) datasets. (A-C) n, number of cells per cluster.

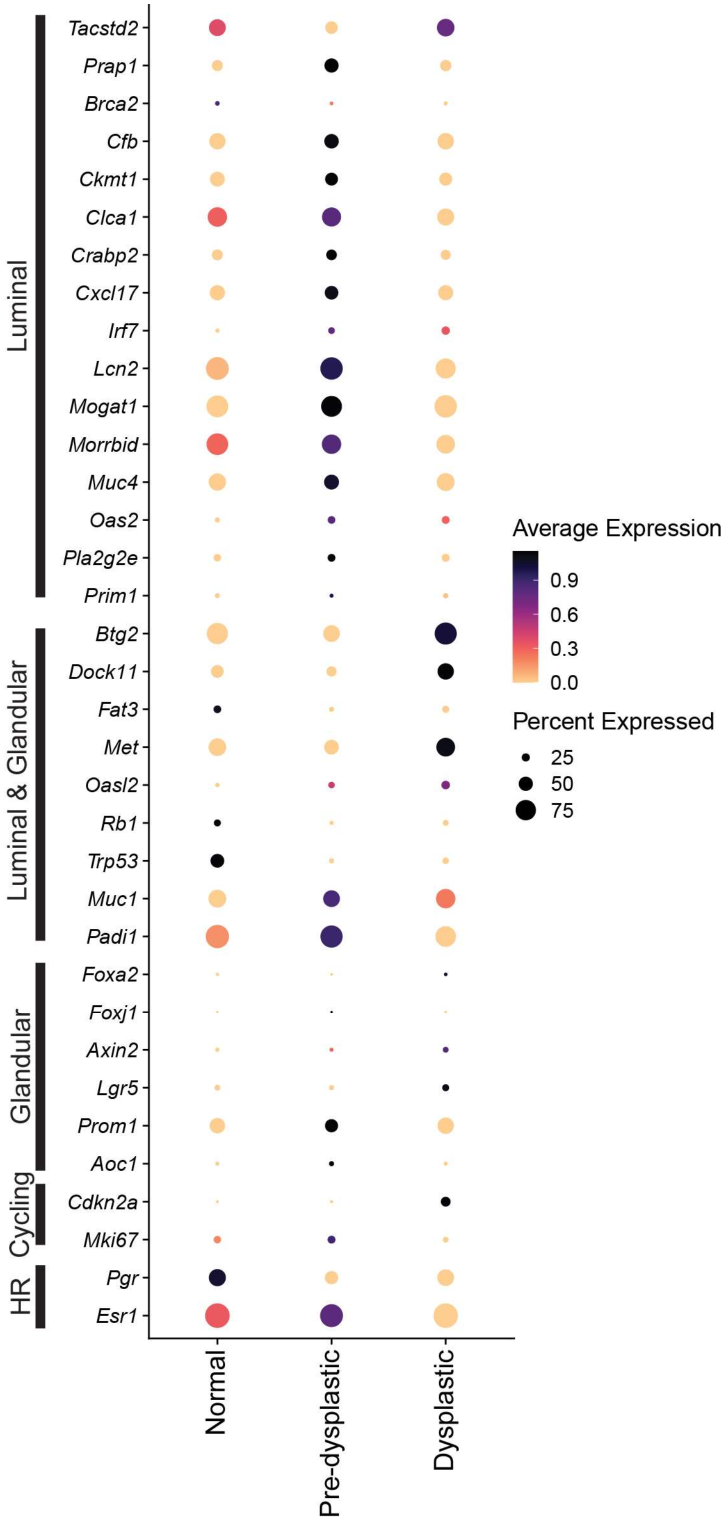


**Supplementary Figure 6. Differentially expressed genes across SEC stages with their spatial location.** A dot plot showing the expression of genes identified to be differentially expressed with high consistency in our single-cell and spatial SEC samples and grouped by spatial location in Visium data. Some genes were also added that are known to be involved with gynecological malignancies. Some genes, such as *Tacstd2* and *Foxa2,* are used as cell-type references. HR, Hormonally Responsive; Mito, Mitochondrial.

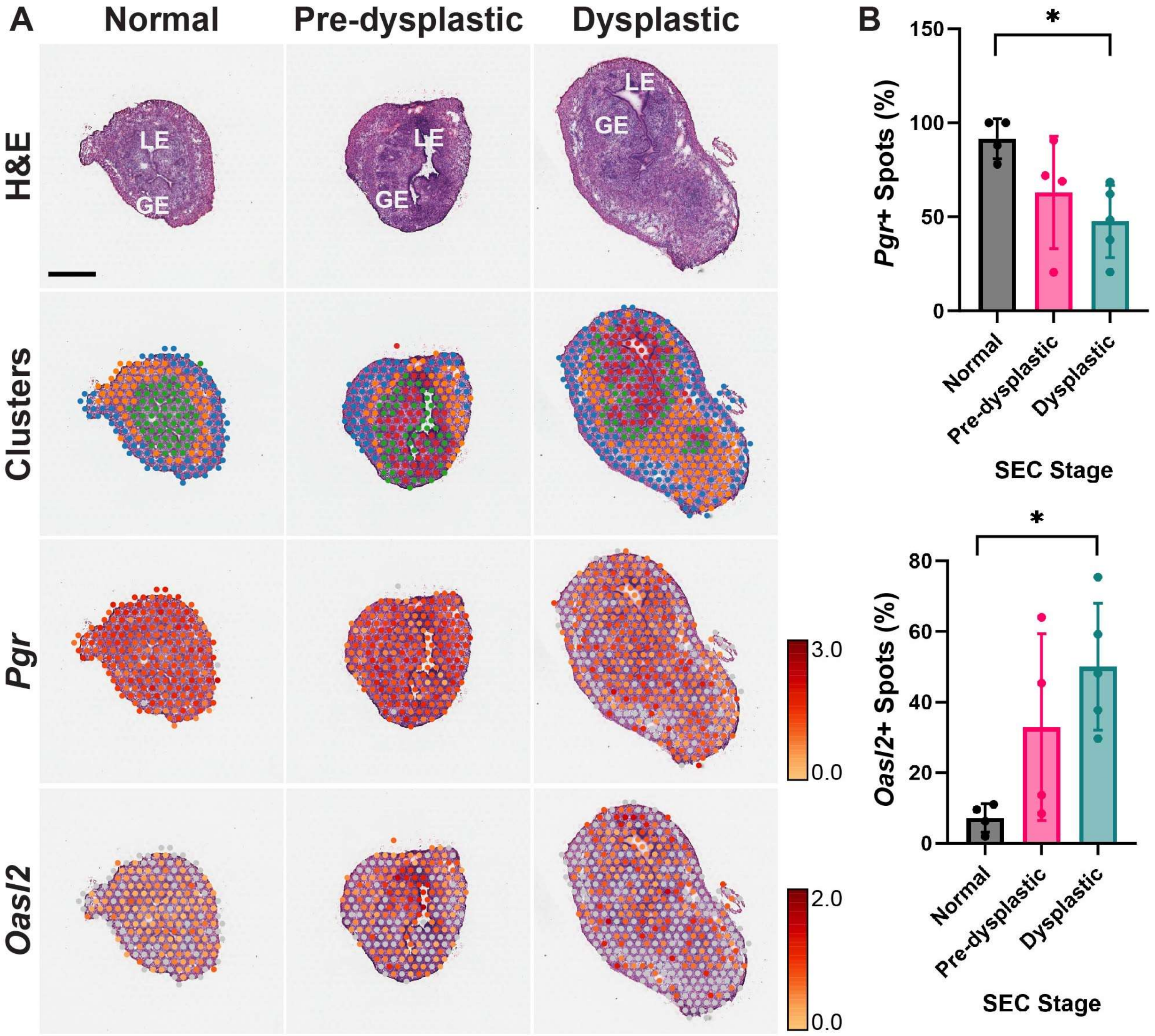


**Supplementary Figure 7. Expression of *Pgr* and *Oasl2* during carcinogenesis.** (A) Histological (Hematoxylin and Eosin, H&E) and spatial transcriptomic (Clusters) visualization, and detection of *Pgr* and *Oasl2* mRNAs in normal, pre-dysplastic, and dysplastic endometrial epithelium. Visium spatial RNA sequencing. Scale represents the log normalized expression of the spots across each individual sample. (B) Quantification of *Pgr*+ (top) and *Oasl2*+ (bottom) spots within the endometrium in normal (n=4), pre-dysplastic (n=4), and dysplastic (n=5) Visium samples. A cutoff of 25% of the maximum log normalized expression value determined spot positivity. (A) Scale, top left, 1 mm. (B) *P<0.05, Mann-Whitney U Tests. All error bars denote s.d.

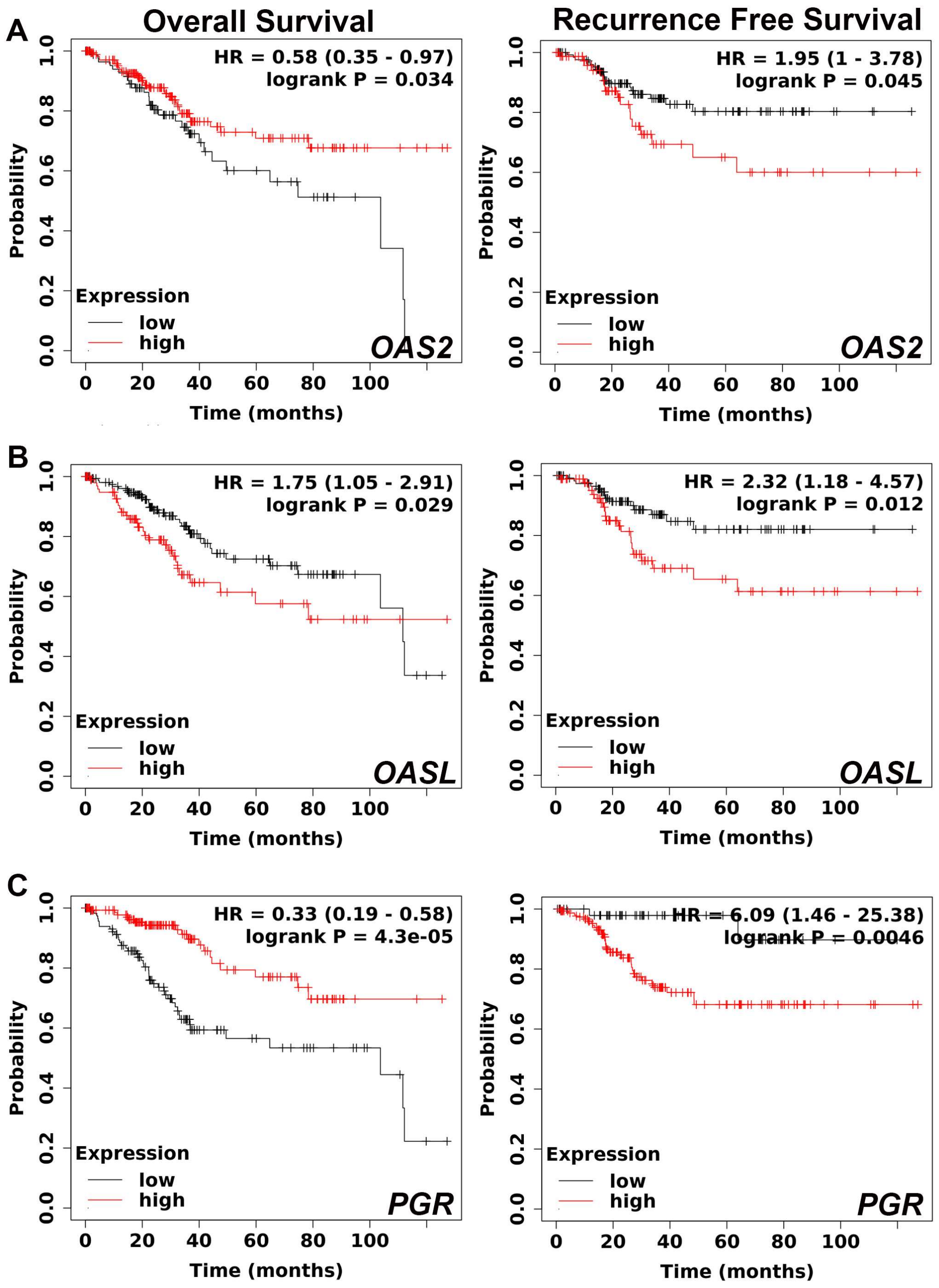


**Supplementary Figure 8. Survival curves for diagnostic genes.** (A-C) Kaplan-Meier plots for mRNA expression of *OAS2* (A), OASL (B) and PGR (C) with overall survival (left) and recurrence free survival (right). (A-C) Survival was stratified using a cutoff of *CDKN2A* expression as a diagnostic marker of SEC, where only cases above the median expression of *CDKN2A* were evaluated. Data collected from KMPlotter[29].

**Supplementary Table 1. Differentially expressed genes confirmed by spatial analysis.**

| Diff Exp Genes | Impact on Patient Survival |
|---|---|
| *Add2* | Yes* |
| *Alox12e* | Yes |
| *Alox15* | Yes |
| *Aoc1* | Yes |
| *Astn2* | Yes |
| *Auts2* | Yes |
| *Bcl2l15* | Yes |
| *C3* | Yes |
| *Cdkn2a* | Yes |
| *Cfb* | Yes |
| *Ckmt1* | Yes |
| *Clca1* | Yes |
| *Crabp2* | Yes |
| *Ctsd* | Yes |
| *Cxcl17* | Yes |
| *Dock11* | Yes |
| *Eps8* | Yes |
| *Epsti1* | Yes |
| *Ern1* | Yes |
| *Fcgbp* | Yes |
| *Gclc* | Yes |
| *Irf7* | Yes |
| *Itgam* | Yes |
| *Kitl* | Yes |
| *Lcn2* | Yes |
| *Lpar3* | Yes |
| *Ltf* | Yes |
| *Ly6a* | Yes |
| *Magi3* | Yes |
| *Met* | Yes |
| *Mogat1* | Yes |
| *Muc1* | Yes |
| *Muc4* | Yes |
| *Oas2* | Yes |
| *Oasl2* | Yes |
| *Padi1* | Yes |
| *Pim1* | Yes |
| *Pla2g2e* | No |
| *Plet1* | Yes |
| *Ppp2r2b* | Yes |
| *Ptprj* | Yes |
| *Rn7sk* | Yes |
| *Scd2* | Yes |
| *Sprr2f* | Yes |
| *Tnfaip8* | Yes |

*Yes, significant decrease in survival.

**Supplementary Table 2. Antibodies used for immunostaining.**

| Antigen, conjugation | Antibody source, catalogue number | Clone | Host | Retrieval, Dilution |
|---|---|---|---|---|
| FOXA2 | Abcam, ab108422 | EPR4466 | Rabbit | Citrate, 1:300 (IF[&]), 1:750 (IP-ABC[#]) |
| Ki67 | Invitrogen, 14-5698-82 | SolA15 | Rat | Citrate, 1:4000 (IP-ABC) |
| OAS2 | Abcam, ab197655 | PC* | Rabbit | Citrate (30 min) 1:400 (IP-ABC) |
| P16 | Abcam, ab241543 | PABLO33B | Rat | Citrate (20 min) 1:500 (IP-ABC) |
| PGR | Abcam, ab63605 | PC | Rabbit | Citrate (15 min) 1:200 (IP-ABC) |
| tdTomato/RFP | Rockland Immunochemicals Inc., 600-401-379S | PC | Rabbit | No retrieval, 1:4000 (IP-ABC) |
| TROP2 | Santa Cruz Biotechnology, sc-376181 | F-5 | Mouse | Citrate (15 min) 1:50 (IF), 1:750 (IP-ABC) |
| Anti-mouse IgG, biotinylated | Vector Labs, BA-9200-1.5 | PC | Goat | 1:200 |
| Anti-mouse IgG, Alexa Fluor 488 | Thermo Fisher, A-21208 | PC | Goat | 1:200 |
| Anti-rabbit IgG, biotinylated | Vector Labs, BA-1000-1.5 | PC | Goat | 1:200 |
| Anti-rabbit IgG, Alexa Fluor 594 | Thermo Fisher, A-21207 | PC | Donkey | 1:200 |
| Anti-rat IgG, biotinylated | Vector Labs, BA-4000-1.5 | PC | Rabbit | 1:200 |

*PC, polyclonal
[#]IP-ABC, immunoperoxidase-ABC Elite method
[&]IF, immunofluorescence
